# Insights into the phenology of migration and survival of a long migrant land bird

**DOI:** 10.1101/028597

**Authors:** Bénédicte Madon, Eric Le Nuz, Cédric Ferlat, Yves Hingrat

## Abstract

**Lay summary:** For polygamous long-migrant birds, the choice of migration strategy depends on social pressure and experience and influences the chance of survival. If you are a male, you’d better leave early in the spring to secure the best site to show off. In fall, juveniles have a hard time surviving to migration as they leave before the adults and lack experience on where to go and where to stop to rest.

**Abstract:** The process of migration stems from an adaptation of climatic seasonality and animals have developed various strategies to complete the journey between a wintering and breeding ground. Understanding the migratory behavior and determining when and where mortality occurs during the annual cycle is fundamental to understand population dynamics and implement appropriate conservation measures. Based on a big data set and advanced statistical methods, we inspected the phenology of migration of a polygynous land bird, the Macqueen’s bustard, *Chlamydotis macqueenii.* We explored its migration strategies between sex, age, season and geographical origin. We show that departure for migration depended on age in the fall with juveniles being the first to leave and on age and sex in the spring with juveniles departing later and males induced to arrive early in spring to secure high-quality territories. Birds breeding at higher latitudes were the first to leave in the fall and more likely to perform longer stopovers. Bustards exhibited different strategies for spring and fall migrations: spring migration was significantly longer than fall migration with more but shorter stopovers. Survival was lower for juveniles experiencing their first migration and for all birds during fall migration and on their wintering ground. Experience linked to social hierarchical pressures and environmental conditions might be the key drivers of migration strategies and survival in long-distance polygynous migrants.

## Introduction

The annual cycle of migratory birds stems from an adaptation to climatic seasonality and is typically composed by three major events of variable timing, duration and sequencing: breeding, molt and the return journey between wintering and breeding grounds, i.e., the migration (Somveille et al. 2015). Different sets of rules determining the process of migration (Alerstam et al. 2006, Duriez et al. 2009), called migration strategy, have been highlighted and three main hypotheses were proposed (for a review see Ketterson and Nolan 1983) to explain the differences in migration strategy between individual classes (e.g., age, sex, reproductive status). The ‘Arrival Time’ hypothesis invokes that the reproductive fitness of one sex is partly influenced by the acquisition of a territory in early spring, and, as high quality territories are limited, the early arrival of individuals is an advantage to secure a territory acquisition (Kokko 1999). The ‘dominance’ hypothesis (or ‘competitive release’ hypothesis) posits that food scarcity drives subordinate individuals to migrate further to limit food competition (Rogers et al. 1989). The ‘body size’ hypothesis (or ‘thermal tolerance’ hypothesis) suggests that thermal efficiency dictates migratory tendency, with smaller individuals being more likely to migrate further. These hypotheses stem from the study of bird migration which has been largely dominated by studies on bird species benefitting from intensive ringing programs (Bairlein 2001). However, for species with no other movement monitoring options (limited field access: ocean and desert crossing species), the development of remote monitoring tools such as satellite tracking brought a much needed salvation and has opened new perspectives (Arizaga et al. 2014). This is the case of the Macqueen’s bustard, *Chlamydotis macqueenii,* a partial migrant bird species (Goriup 1997), classified as Vulnerable (BirdLife International 2014). From the early 90’s, an intensive monitoring of migrant individuals using satellite tracking was launched by the National Avian Research Centre (Abu Dhabi, United Arab Emirates) and laid the foundations for the early study of the species. Migrant populations were shown to breed from west Kazakhstan to China and winter in the range of resident populations in South Central Asia and the Middle-East (Combreau et al. 2001, Combreau et al. 2011b). On their breeding ground, migrant Macqueen’s bustard exhibit a polygynous mating system where males compete for display territories to which they remain faithful during the breeding season (Riou and Combreau 2014). This monitoring effort has been reinforced to this day, with more than 400 birds equipped with satellite transmitters in central Asia. This unprecedented data set offers the opportunity to better highlight the migration strategies among sex and age-classes in a rarely-studied system, i.e. a polygynous land bird (but see Kessler et al. 2013, Garcia De La Morena 2015), in the light of the three main hypotheses: arrival-time, dominance and body-size. Site fidelity and intra-sexual competition are likely to be the main drivers for male migration timing and distance (Schroeder and Robb 2003; Boyle 2008), suggesting the ‘arrival time’ hypothesis. Females and juveniles, whose fitness depend less on securing a breeding site and whose survival might be influenced by their smaller size (Martín et al. 2007), might have an obligate strategy due to social hierarchical pressures of male dominance, suggesting the ‘dominance’ and ‘body size’ hypotheses.

The chosen migration strategy will likely influence the annual survival of individuals, which is the product of survival rates at the four periods of their annual cycle: breeding, fall migration, wintering, and spring migration. Tracking data sets can be converted in capture-recapture histories allowing advanced survival analyses (Duriez et al. 2009, Hardouin et al. 2014) taking into account such temporal breakdown. However, the extent of differential migratory patterns and their relation to differential survival rates has rarely been explored (Hutto 2000, Sillett and Holmes 2002, Lok 2011). Yet understanding the migratory behavior and determining when and where mortality occurs during the annual cycle is fundamental to understand population dynamics and implement appropriate conservation measures (Leyrer et al. 2013, Klaassen et al. 2014).

Based on an eight-year satellite-tracking data set, we inspected the full picture of migration and survival of the Macqueen’s Bustard. Using recent advances in movement analyses, we were first able to determine individual movement key timings. Then, using robust statistical analyses and multistate capture-recapture modelling, we highlighted the influence of individual traits (age and sex) and spatio-temporal factors (geographic origin and season) on the migration strategy and survival of a polygynous long-migrant land bird species.

## Materials and methods

### DATA

A total of 414 wild migrant Macqueen’s Bustards were captured during the breeding season (end of March to end of June) between 2010 and 2013 in Uzbekistan (Navoi district, 39°N, 65°E) and between 2005 and 2013 in Kazakhstan (Central Kazakhstan: Shimkent area, 43°N, 67°E; East Kazakhstan: 46°N, 78°E; West Kazakhstan: Mangystau area, 42.75°N, 52°E; Fig.1). Adult birds were trapped using loop cord snares. Males were trapped on their display sites baited by a dummy female and females were trapped on their nest replacing live eggs with wooden eggs (see method in Hardouin et al. 2014). Juveniles were trapped by hand before fledging (see method in Combreau et al. 2002 and Hardouin et al. 2011). All birds were weighted and ringed. Males weighted on average 2 kg ±1.7, females 1.3 kg ±1.3 and juveniles 0.7 kg ±1.3. Birds were equipped GPS-PTT (platform terminal transmitter) solar-powered satellite transmitters (Microwave Telemetry Inc, Columbia, MD, USA) of 22 to 45g depending on bird weight (representing on average 3%±1 of individuals weight (Kenward 2001). Transmitters were operated through the ARGOS system in Toulouse (CLS, France) and programmed to record a GPS position every two hours and transmit once every two days.

**Fig. 1.**
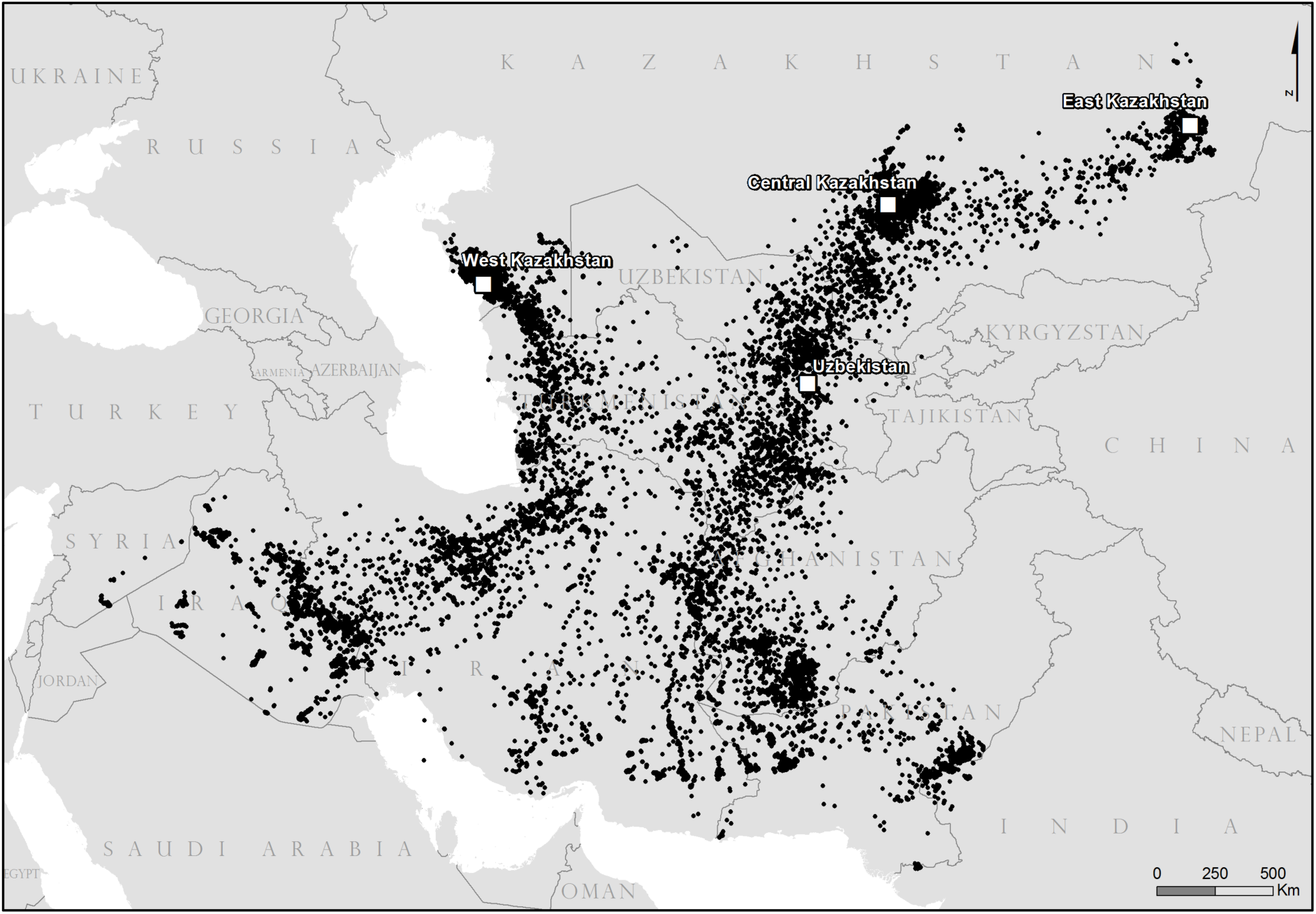
Places of origin (trapping locations) of Macqueen’s bustards equipped in Uzbekistan and in Kazakhstan (East, Central and West Kazakhstan) between 2005 and 2013. Black dots are daily locations of 150 wild adults and 51 wild juveniles retained for analyses.

Satellite tracking data from birds that did not migrate before the transmitter stopped transmitting or with missing data (See Madon and Hingrat 2014) were not included in the analysis. Hence a total of 158 wild adults and 41 wild juveniles were included in the analyses (Table 1). Data were first filtered by precision: GPS and ARGOS locations of CLS classes 2 and 3 were selected. The last daily location was then retained for each individual to allow for regular time spacing (i.e., an approximate 24h gap) between successive locations. Location coordinates were then projected using the Asia north equidistant conic projection in ArcGIS 10.1 (ESRI 2012) to calculate distances between successive locations (in km), i.e., daily distances, and build a daily distance time series for each bird.

**Table 1.** Sample sizes by sex and geographical areas between 2005 and 2013 of PTT-equipped equipped individuals, subset of individuals included in the analyses with their monitoring lengths (and associated standard error) in terms of complete migration numbers, and timing of fall and spring migration as median departure dates (day of the year) with associated standard error.

| Country | Region | Sex | Number in |  |  |  |  |
| --- | --- | --- | --- | --- | --- | --- | --- |
|  |  |  | Individuals<br>equipped | the<br>analyses | Monitoring<br>length | Mean fall<br>date | Mean spring<br>date |
| Kazakhstan | West | female | 43 | 37 | 3.1 (0.5) | 14 oct (2.5) | 6 mar (1.4) |
|  |  | male | 19 | 19 | 2.5 (0.5) | 19 oct (4) | 26 feb (2.1) |
|  |  | Juvenile | 23 | 19 | 1.5 (0.4) | 30 sept (2.7) | 24 mar (2.1) |
|  | Central | female | 213 | 47 | 3.3 (0.4) | 18 oct (1.6) | 11 mar (1.5) |
|  |  | male | 30 | 11 | 2.4 (0.4) | 7 nov (3.6) | 2 mar (2.7) |
|  |  | Juvenile | 34 | 26 | 0.6 (0.2) | 18 oct (2.8) | 1 ap (7.2) |
|  | East | female | 20 | 18 | 2.8 (0.5) | 10 sept (2.6) | 23 feb (3.5) |
|  |  | male | 9 | 9 | 2.8 (0.8) | 22 sept (4.9) | 18 feb (3.2) |
|  |  | Juvenile | 8 | 6 | 0.3 (0.3) | 12 sept (4.6) | 12 ap (na) |
| Uzbekistan | Navoi | female | 13 | 6 | 2.5 (0.6) | 17 oct (5.2) | 4 mar (3.7) |
|  |  | Male | 4 | 3 | 1.3 (1.3) | 8 nov (3.4) | 24 feb (16.5) |

The PELT-TREE method was used to break down the daily distance time series of each bird. This recent framework combines a change point algorithm to find the changes in variance, the so called change points, in the daily distance series for each bird and a classification tree to classify the obtained segments. Here we considered three movement behavioral classes: staging, migratory and non-migratory movements. Based on the training data used by the classification tree, the mathematical rules to classify the segments into the three movement classes were defined as follow: segments with mean < 17.642 km were classified as “staging”, segments with mean >17.642 km and < 100.284 km as “non-migratory” and segments with mean > 100.284 km as “migratory” (See Madon and Hingrat 2014 for details).

Based on the segmentation, we defined, for each bird, key timings of migration as follow: departure date, i.e., start of migration, as the first day of migratory movement (or non-migratory movement if immediately followed by a migratory movement) following a staging period, in the opposite direction compared to the preceding migratory movement; arrival date, i.e., end of migration, as the first day of staging after a migratory movement (or non-migratory movement if immediately following a migratory movement), given that the next migratory movement is in the opposite direction; and stopover as any segment of staging behavior between the departure and arrival dates.

### STATISTICAL MODELLING

#### Migration strategy

We explored the fall and spring migration strategy of the Macqueen’s bustard in terms of six response variables: 1- fall and spring migration departure dates, 2- migration distances, i.e., sum of the daily distances (in km) between the migration departure and arrival dates, 3- migration duration, i.e., number of days between the departure and arrival dates, 4- number of stopovers, i.e., number of staging segments between the migration departure and arrival dates.

The variable migration duration was further broken down into two variables in the analyses: 5- duration of migratory movement, i.e., total length in days of migratory and non-migratory movement segments between the migration departure and arrival dates, 6- duration of stopovers (Alerstam et al. 2006), i.e., total length in days of staging segments between the migration departure and arrival dates.

We conducted two sets of linear (or generalized linear) mixed model (See (Bolker et al. 2008) for a review) analyses on each response variable with individual and year as random factors (Table 2) using four data sets: dataset 1- all individuals, dataset 2- sexed individuals, i.e., adults only, dataset 3- all individuals presenting at least 1 stopover, and dataset 4- sexed individuals (adults only) presenting at least one stopover (Table 2).

**Table 2.** Modeling approaches (model type: lqmm = linear quantile mixed model on median; lmm = linear mixed model glmm = generalized linear mixed model; data distribution: “Family” (with the link for the glmm); R package and function) for the following migration response variables: fall and spring departure date, migration distance, duration of movement, number and duration of stopovers, using 4 datasets (“1” = all individuals, “2”= sexed individuals, i.e., adults only, “3” = all individuals presenting at least 1 stopover, “4” = sexed individuals presenting at least 1 stopover) and factor “Age”, “Place”(i.e., breeding place), “Sex” (adult birds) and “Season” as explanatory fixed factors and individual and year as random factors.

| Explanatory<br>Response | Explanatory |  |  |  | Model type | Family (link) | R Package: function |
| --- | --- | --- | --- | --- | --- | --- | --- |
|  | Age | Place | Sex | Season |  |  |  |
| Fall departure date<br>Spring departure date | 1 | 1, 2 | 2 |  | lqmm | Normal | lqmm: lqmm() |
| Migration distance | 1 | 1,2 | 2 | 1,2 | lmm | Normal | lme4: lmer() |
| Migration duration | 1 | 1,2 | 2 | 1,2 | glmm | Poisson (log) | lme4: glmer() |
| Duration of movement | 1 | 1,2 | 2 | 1,2 | glmm | Poisson (log) | lme4: glmer() |
| Number of stopovers | 1 | 1,2 | 2 | 1,2 | glmm | Binomial (logit)/ | lme4: glmer() |
|  | 3 | 3,4 | 4 | 3,4 |  | Poisson (log),<br>Poisson (log) |  |
| Duration of stopovers | 3 | 3,4 | 4 | 3,4 | glmm | Poisson (log) | lme4: glmer() |

With datasets 1 and 3, we used explanatory fixed factors “Age” (available for all individuals), “Place” (corresponding to the breeding place for the adults and birth place for the juveniles), “Season” (except for response variables fall and spring departure dates) to model the response variables. Because Macqueen’s bustards may start breeding from one year old (Saint Jalme and van Heezik 1996), only the first year of monitoring of juveniles was included in the analyses to account for them as non-breeders. So factor “Age” refers to the reproductive status of individuals. With datasets 2 and 4, we used factor “Sex”, along with factors “Place” and “Season” (except for response variables fall and spring departure dates)(Table 2). Only the interaction Sex*Season was considered, due to small sample sizes in levels of other interactions and factors were considered significant when *p* < 0.05 or |*t*| > 2 (*p* being unavailable in package ‘lmm’) (Baayen et al. 2008, Bolker et al. 2008). All analyses were conducted in R (R Core Team 2014).

#### Survival

We used multistate capture-recapture models to estimate survival by describing the transition between the states “alive” and “dead” (Lebreton et al. 1992). These models are defined in terms of three processes (initial state, event and state processes) allowing the simultaneous estimation of: the encounter probability (the probability that an individual is encountered in site *A* and time *t* given that it is alive in site *A* and time *t*), the apparent survival (the probability that an individual alive at site *A* and time *t* is still alive at time *t*+1) and transition between sites, i.e., movements (the probability that an individual moves from site *S* at time *t* to site *Z* at time *t*+1, given that it survived from time *t* to *t*+1; hence denoted “transition matrix” or “movement probabilities” conditional on survival) (Lebreton and Pradel 2002). Here we dealt with a mixture of live recaptures and dead recoveries (e.g., Duriez et al. 2009, Le Gouar et al. 2011), reported when the transmitters were retrieved in the field. Hence survival was modelled as a transition from the state “alive” to the state “newly dead”. Encounter histories were split in four yearly occasions, each corresponding to one “movement phase” with two fates, i.e., alive recaptures and dead recoveries. The four occasions corresponded to the four seasonal phases of movement of a migratory animal determined by the above key timings: on the breeding ground (period between the spring arrival date and fall departure date), in fall migration (period between the fall departure date and the fall arrival date), on the wintering ground (period between the fall arrival date and the spring departure date) and in spring migration (period between the spring departure date and the spring arrival date). We thus accounted for nine states: four alive states (1-4) and four newly dead states (5-8) in the seasonal phases of movement and one unobserved dead state (9). Given that individuals were equipped with GPS-PTT transmitter, the successive states occupied by an individual can be observed directly and the encounter probability in the four alive states is consequently equal to 1. In the transition matrix, movement and survival are considered as two successive steps. Here, if a bird was found dead during a movement phase, it had necessarily moved from the previous movement phase before dying. Therefore, movements were estimated before survival in the transition matrix, i.e., the survival probability depends on the site of arrival, e.g., in Duriez et al. (2009).

Difficulties in attributing precisely the “movement phase” arose when a bird died after starting migration, as it was not possible to determine whether it was still migrating or had arrived on the wintering/breeding site before dying. Thus, we considered that an individual was newly dead on the breeding ground: 1- when transmitters were retrieved on the breeding ground or 2- when the individual was lost after the 1^st^ of July (i.e., the signal was suddenly lost or it was reported non-moving with the same position before loss of the signal but the transmitter was not retrieved in the field). Similarly we considered that an individual was newly dead on the wintering ground: 1- when PTT transmitters were retrieved on the wintering ground or 2- when the individual was lost after the 1^st^ of January.

Each step of a multistate model, i.e., initial state, event process and state process, can be parametrized with environmental covariates or individual factors. Here we focused on individual factors “Experience”, “Sex*Age” (males, females and juveniles), “Place” (Central Kazakhstan, East Kazakhstan, West Kazakhstan and Uzbekistan) and time factors. Factor “Experience” was related to age at capture and consisted in two groups: “first timers” and “experienced birds”. The group “first timers” included the first year of monitoring of birds equipped as juveniles on the breeding ground, hence first timers in terms of fall migration, wintering and following spring migration. The group of “experienced birds” corresponds to birds equipped as adults and to juveniles after a first year of monitoring (from their second spring after their first fall migration). Time factor included “4 periods”: time divided in the four movement phases. We also tested time divided into “2 periods” with time periods pulled into 2 main periods “spring migration and breeding ground” and “fall migration and wintering ground”, to account for the difficulties in attributing death to these successive periods.

Model selection was performed using program E-SURGE v1.8.9 (Choquet et al. 2009) with an Akaike Information Criterion corrected for sample size calculated as follows: QAICc = (deviance/*ĉ*) + 2*K* + (2*K*(*K*+1))/(*n*-*K*-1), where *K* and *n* are the number of parameters and the effective sample size respectively. The preferred model was the one with the smaller QAICc value and two models were deemed to be equivalent when they differed by less than two. In addition to the QAICc, we paid attention also to the biological plausibility and quality (confidence intervals) of the estimates when selecting models. We used a generalized logit-link function. Description of the model structure and matrix patterns used in the models developed in E-SURGE is given in Appendix S1.

## Results

Among Macqueen’s bustards equipped in Central Asia with GPS-PTT transmitter between 2005 and 2013, we obtained accurate data for our analysis from 201 birds (Table 1). Birds were followed on average over two migrations (*se =* 0.07), i.e one year. Two females from West Kazakhstan were followed during 10 and 14 migrations, i.e., five and seven years.

### MIGRATION JOURNEY CHARACTERISTICS

#### Sex-based differential migration

Among adults, there was no difference in the timing of departure between males and females in the fall but in the spring, males departed for migration 8 (*se* = 2.98, lqmm *p* = 0.01) days earlier (Table 1). Migration distance was similar and both sexes were as likely to perform stopovers. Migration duration, which included stopovers and movements, was significantly shorter for males (glmm (logit scale) *β* = −0.24 (*se* = 0.1), *p* = 0.02) (Table 3). There was no difference in terms of duration of movement but the time spent on stopovers by males was significantly shorter (glmm (log scale) *β* = −0.3 (*se* = 0.12), *p* = 0.01).

**Table 3.**
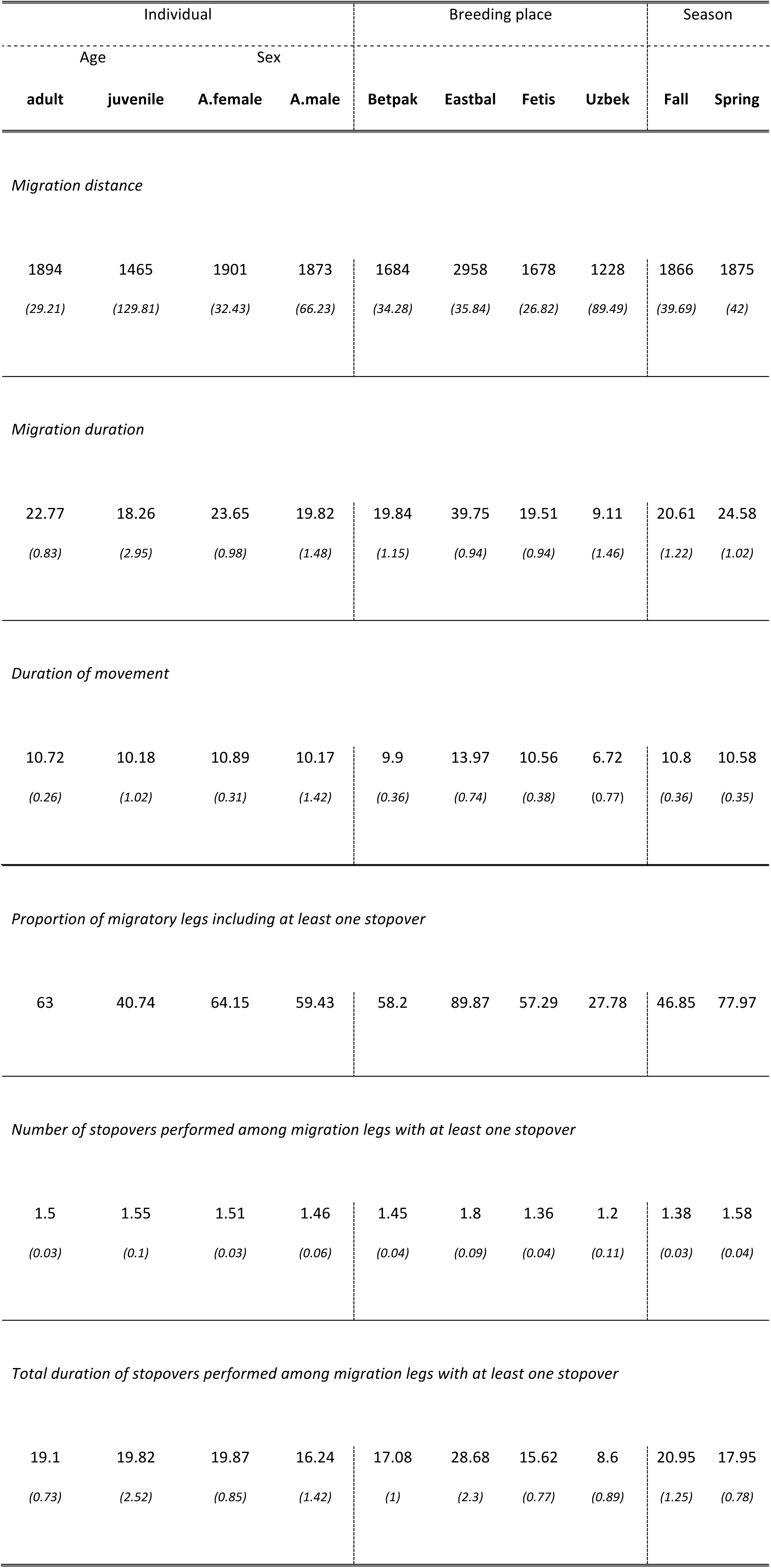
Mean (*and standard error*) *of* migratory distances (in km), migration duration (in days), duration of movement (in days) and proportion (%) of migratory legs with at least one stopover and among those, mean (*and standard error*) total duration (in days) and number of stopovers for the fall and spring migrations of wild Macqueen’s bustards breeding in Asia (in Uzbekistan (Uzbek) and Kazakhstan: Betpakdalah (Betpak), Eastbalkash (Eatbal), Fetisovo (Fetis)), and equipped with PTT transmitter between 2005 and 2013.

| Individual |  |  |  | Breeding place |  |  |  | Season |  |
| --- | --- | --- | --- | --- | --- | --- | --- | --- | --- |
| Age |  | Sex |  |  |  |  |  |  |  |
| adult | juvenile | A.female | A.male | Betpak | Eastbal | Fetis | Uzbek | Fall | Spring |
| *Migration distance* |  |  |  |  |  |  |  |  |  |
| 1894 | 1465 | 1901 | 1873 | 1684 | 2958 | 1678 | 1228 | 1866 | 1875 |
| (29.21) | (129.81) | (32.43) | (66.23) | (34.28) | (35.84) | (26.82) | (89.49) | (39.69) | (42) |
| *Migration duration* |  |  |  |  |  |  |  |  |  |
| 22.77 | 18.26 | 23.65 | 19.82 | 19.84 | 39.75 | 19.51 | 9.11 | 20.61 | 24.58 |
| (0.83) | (2.95) | (0.98) | (1.48) | (1.15) | (0.94) | (0.94) | (1.46) | (1.22) | (1.02) |
| *Duration of movement* |  |  |  |  |  |  |  |  |  |
| 10.72 | 10.18 | 10.89 | 10.17 | 9.9 | 13.97 | 10.56 | 6.72 | 10.8 | 10.58 |
| (0.26) | (1.02) | (0.31) | (1.42) | (0.36) | (0.74) | (0.38) | (0.77) | (0.36) | (0.35) |
| *Proportion of migratory legs including at least one stopover* |  |  |  |  |  |  |  |  |  |
| 63 | 40.74 | 64.15 | 59.43 | 58.2 | 89.87 | 57.29 | 27.78 | 46.85 | 77.97 |
| *Number of stopovers performed among migration legs with at least one stopover* |  |  |  |  |  |  |  |  |  |
| 1.5 | 1.55 | 1.51 | 1.46 | 1.45 | 1.8 | 1.36 | 1.2 | 1.38 | 1.58 |
| (0.03) | (0.1) | (0.03) | (0.06) | (0.04) | (0.09) | (0.04) | (0.11) | (0.03) | (0.04) |
| *Total duration of stopovers performed among migration legs with at least one stopover* |  |  |  |  |  |  |  |  |  |
| 19.1 | 19.82 | 19.87 | 16.24 | 17.08 | 28.68 | 15.62 | 8.6 | 20.95 | 17.95 |
| (0.73) | (2.52) | (0.85) | (1.42) | (1) | (2.3) | (0.77) | (0.89) | (1.25) | (0.78) |

#### Age-based differential migration

Juveniles departed for migration significantly earlier than adults in fall (6 days, *se* = 2.79, lqmm *p* = 0.03) and later in spring (27 days, *se* = 3.37, lqmm *p* <0.05) (Table 1). On average, juveniles travelled as long as adults for both migrations in terms of distance and migration duration and were as likely to perform stopovers. However, time spent on stopover was significantly longer (glmm (log scale) *β* = 0.36 (*se* = 0.17), *p* = 0.03)(Table 3).

#### Geographical origin-based differential migration

Birds departing from East Kazakhstan, migrated more than twice as far as birds from lower latitudes (i.e., on average 2958 (*se* = 35.84) km for East Kazakhstan against 1228 (*se* = 84.65) km for Uzbekistan) (Fig. 1, Table 3). As a consequence the median dates of departure followed a latitudinal gradient for both the fall and spring migrations: birds from breeding grounds at lower latitudes left later for migration. For the fall migration, the median date departure of birds from higher latitudes departed significantly earlier. Compared to birds from Central Kazakhstan, birds from East Kazakhstan departed 37 (*se* = 3.44.21, lqmm *p* <0.05) days earlier and birds from West Kazakhstan 9 (*se* = 2.43.55, lqmm *p* <0.05) days earlier. Birds from Central Kazakhstan departed slightly earlier (4 days) than birds from Uzbekistan (Table 1). In the spring, the departure dates did not follow the latitudinal gradient. Adult birds from Central Kazakhstan were the last ones to depart for migration on 8 March (67^th^ day, *se* = 1.43): West Kazakhstan birds departed 6 days earlier (*se* = 2, lqmm *p* < 0.05), Uzbekistan birds 7 days earlier (*se* = 2.9, lqmm *p* = 0.02), and East Kazakhstan birds 14 days earlier (*se* = 4.25, lqmm *p* <0.05).

Birds with longer migratory journeys were more likely to perform stop-overs (Table 3). For example, birds from East Kazakhstan which travelled on average 1328 km (*se* = 92.05, lmm *t* = 14.43) more than birds from Central Kazakhstan, were 10 times more likely to perform stopovers (glmm (logit scale) *β* = 2.3 (*se* = 0.53), *p* < 0.05), 90% of their migration legs displayed at least one stopover and they spent significantly longer periods at stopover sites (on average 28.68 (*se* = *2.44*) days, glmm (log scale) *β* = 0.63 (*se* = 0.15), *p* < 0.05). On the other hand, birds from Uzbekistan were less likely to stop for refueling (glmm (logit scale) *β* = − 1.66 (*se* = 0.75), *p* = 0.03) and only 28% of their migration legs included a stopover (Table 3).

#### Season-based differential migration

Macqueen’s bustards appeared to exhibit different behaviors in spring and fall migrations. Results indicated that spring migration was significantly longer than fall migration (respectively 24.52 (*se* = 1.03) days against 20.66 (*se* = 1.24) days, glmm (log scale) *β* = 0.24 (*se* = 0.023), *p* < 0.05) (Table 3). However, spring migration was significantly shorter in terms of movement duration for males (interaction sex*season, glmm (log scale) *β* = −0.17 (*se* = 0.07), *p* = 0.015)). In terms of refueling strategy, birds were more likely to stop during the spring migration (glmm (logit scale) *β* = 1.8 (*se* = 0.26), *p* < 0.05) and performed more stopovers in the spring (glmm (log scale) *β* = 0.66 (*se* = 0.099), *p* < 0.05). However, time spent on stopovers was longer during fall migration (glmm (log scale) *β* = −0.07 (*se* = 0.04), *p* = 0.058).

### SURVIVAL

The best fitting model for survival was the model including the interaction of “experience” and the factor “4 periods” where time was divided in the four movement phases (Table 4). First-timers, i.e. juveniles during their first year, had a lower probability to survive at each time period. These differences in survival were especially apparent during their first fall migration (0.62 *se* = 0.07 compared to experienced birds: 0.87 *se* = 0.019) and wintering period (0.65 *se* = 0.086 compared to experienced birds: 0.89 *se* = 0.019). There were no sex-biased mortality patterns (Table 5). Finally, the different migration strategies in fall and spring appeared to impact survival with significantly higher probabilities of surviving the spring migration for both first-timers and experienced birds (respectively 0.9 *se* = 0.067 and 0.97 *se* = 0.01). Survival probabilities were also higher on the breeding ground for experienced birds than on wintering grounds (respectively 0.96 *se* = 0.009 and 0.88 *se* = 0.01).

**Table 4.**
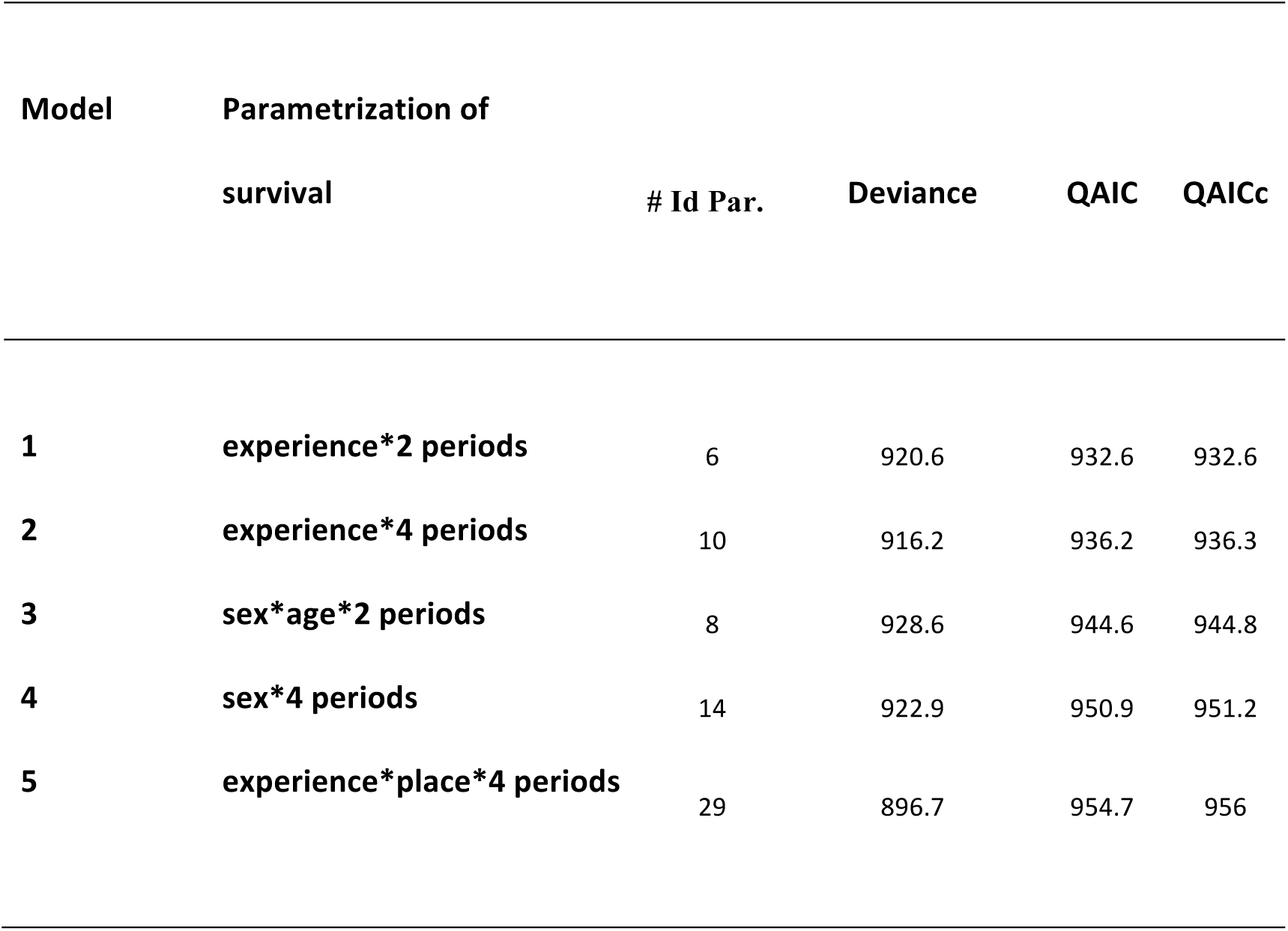
Best models for Macqueen’s bustard survival selected with E-SURGE with associated number of identifiable parameters (# Id Par.), deviance, QAIC and QAICc.

**Table 5.**
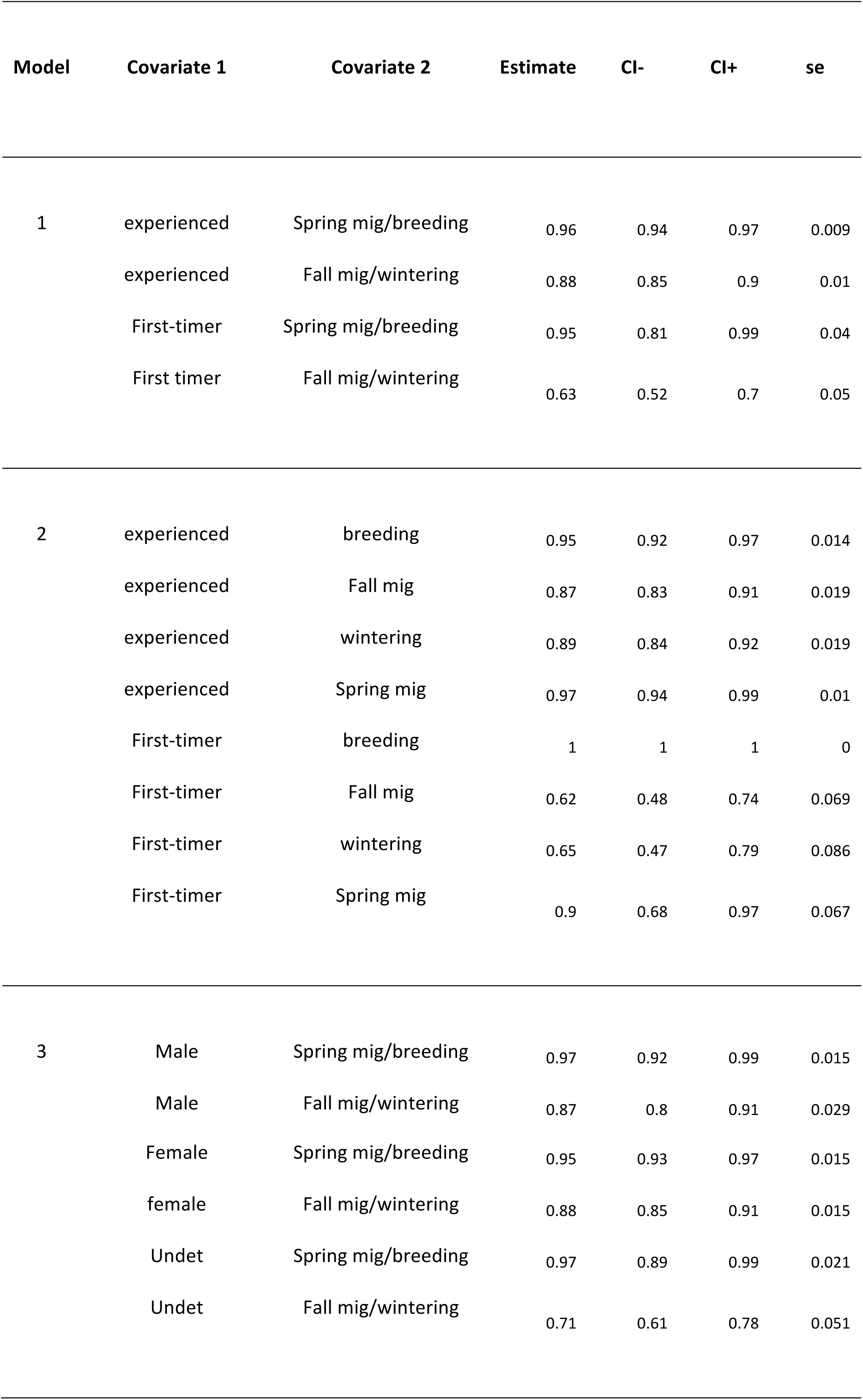
Estimates of survival with associated standard errors (se) and CIs (lower bound: “CI-“, upper bound: “CI+”) for Macqueen’s bustard with the 3 best models selected (Table 4) by E-SURGE. Note that the survival for first-timers is 1 (model 2) for the breeding ground as they have necessarily survived to be included and do not have a following breeding period as they are followed only the first year.

## Discussion

### Timing of migration and survival

The co-existence of different migratory strategies between age and sex groups has been largely discussed and linked to constraints and selective forces in relation to reproductive success, survival and competition. In polygynous species, male competition for display territories during the breeding season is likely to be the main driver for male migration timing (Schroeder and Robb 2003). Our results support the ‘arrival time’ hypothesis: in order to optimize their fitness, Macqueen’s bustard males are induced to arrive early in spring to acquire high-quality territories. Such fitness benefit probably out-weights the cost of migrating out the optimal temporal window, e.g., challenging conditions encountered during late winter-early spring migration (Kokko 1999). Females, on the other hand, can arrive later in spring without reducing their fitness (Kokko et al. 2006). This intersexual out of sync migration timing, e.g., protandry in the spring (Schmaljohann et al. 2015), also warrants females lesser intersexual competition for resources at stopover sites. Interestingly, these differential migration timings do not lead to differences in survival between sexes although for most bird species survival is thought to be higher for males (Sillett and Holmes 2002).

Different factors, such as experience and body condition, are likely to influence migration strategy and survival probability. The body size hypothesis assumes that smaller individuals are less likely to withstand cold temperatures and to experience greater risks associated with fasting in winter (Boyle 2008). This is corroborated by our results showing that, in fall, juveniles leave breeding grounds earlier than adults. Juveniles might be constrained to leave the breeding ground when food and environmental conditions deteriorate because of competition, reduced foraging ability and site-familiarity (Bai and Schmidt 2012). They may be physiologically less capable of undertaking full migration, e.g., different molt process reducing juveniles flying abilities (Newton 2011) or of selecting optimal flight altitude (Mateos-Rodríguez and Liechti 2012). In the Macqueen’s bustard, juveniles which depart earlier do not benefit from social cues to initiate their first fall migration and they cannot use social learning by following adults to locate suitable stopovers and wintering sites and minimize predation risk (Nocera and Ratcliffe 2010, Cresswell 2014). As a consequence, juveniles spent more time on stopovers and had lower survival probabilities during migration as well as on wintering grounds. Greater first-year stochasticity in route-finding, suggesting a bet-hedging strategy (Reilly and Reilly 2009), should nonetheless provide populations of Macqueen’s bustards with greater resilience abilities to large-scale changes (Cresswell 2014). In the following spring, juveniles, which are probably less driven by a breeding pressure, departed later than adults. Little is known about juvenile reproduction timing in Macqueen’s bustards. Studies on North African Houbara bustards, *Chlamydotis undulata undulata,* showed that females initiated reproduction at 1.6 (standard deviation = 0.5) and males at 2.1 (standard deviation = 0.8) years-old (Hardouin et al. 2014). If the pattern of age at first reproduction is similar in the Macqueen’s bustard, it is likely that juvenile migration phenology will be highly variable for the first 2 years (Combreau et al. 2011) and likely more related to natal dispersal (Hardouin et al. 2012). By differing their departure from wintering ground, they might also be able to optimize their survival, hence the high observed survival in the spring migration, by reducing food competition with adults but also by benefiting from the experience acquired in their first migration leg in fall and on wintering ground (Cresswell 2014).

### Refueling and survival

With the development of bird tracking, it has been shown that many species use stopovers along their annual migratory cycle (Guilford et al. 2009, Chevallier et al. 2011, Åkesson et al. 2012). Under the concept of optimal migration, rules for refueling decision at stopover sites have been developed to determine the number of stopovers and time spent on stopovers in order to optimize migration in a given set of constraints (Weber et al. 1999, Duriez et al. 2009, Alerstam 2011). Surprisingly, very few studies have highlighted differential stopover strategies between age, sex, season and geographical origin (Ellegren 1991, Dierschke et al. 2005, Alerstam et al. 2006) and our results demonstrated an effect of each of these factors. As expected, juveniles, that were inexperienced for their first migration, used longer stopovers, a result of different factors detailed above. Our results also highlighted a difference between males and females in terms of time spent on stopovers, with significantly shorter refueling periods for males. This suggests that males use a riskier strategy in spring with faster travel and shorter refueling times in order to optimize their arrival time (Åkesson et al. 2012) or that males have a higher refueling rate (Seewagen et al. 2013). However, these different strategies do not lead to differential survival between sexes. Seasonal differences in migration stopover patterns are also apparent, with individuals performing less but longer stopovers in the fall (Kokko 1999, Alerstam 2006). The longer stopover duration during the fall migration, also observed in some raptor species (Klaassen et al. 2014), might suggest that individuals molt during their fall stopovers and therefore that the role of fall stopovers is twofold: refueling and molting (Hutto 2000). In the case of the Macqueen’s bustard, molting occurs in summer (between end of breeding and migration departure, Gubin 2008) and should not impact the species stopover strategy. Central Asian steppes are characterized by a high productivity during spring which rapidly decreases after summer (Eisfelder et al. 2014). On-route environmental conditions (food limitation and cold temperatures) might be the main drivers for longer stopovers in the fall (Alerstam 2006). In addition, bird condition might be affected by a potential “carry-over effect” of the breeding season (display investments for the males and the parental cares that drain energy reserves for the females). This could explain the observed greater mortality during the fall migration and wintering period, which could be exacerbated by uncontrolled hunting and poaching pressures (Combreau et al. 2001, Combreau 2007). On the other hand, the spring strategy involving short flights interspersed with fewer stopovers to load small fuel reserves assumes that birds will stop at all suitable sites along the migration route making migrants dependent on a chain of sites and consequently more vulnerable to environmental changes in the spring. Under the principle of “multiple jeopardy”, i.e., the probability that any one site is affected by environmental change increases with the number of sites (Newton 2004), birds from higher latitudes (East and West Kazakhstan) which cover greater migratory distances and rely on multiple stopovers (Navedo et al. 2010), will be under greater threat from environmental changes and will be consequently more likely to show declines.

## Conclusion

Little is known about the phenology of migration in polygynous land migrant bird species. Our study provides the first direct evidence of complex migration behaviors and survival: seasonal survival and migration strategies varying by sex, age, season and geographic origin linked to social, hierarchical and physiological pressures. Since direct observations are not possible yet on most parts of the migratory path and wintering ground, we have to rely solely on remote tools and we demonstrate that technology coupled with robust statistical analyses clearly shed light on migration strategies, a key element to implement appropriate conservation measures. Mortality of both adults and juveniles occurs predominantly during the fall migration and the wintering period, similarly to the migrating red knot *Calidris canutus canutus* (Leyrer et al. 2013) and seems to be the driver of decline in many migratory birds (Rappole and McDonald 1994, Carrete et al. 2013). Understanding the relative importance of factors leading to the low survival rates observed during the fall migration and winter (habitat quality versus anthropogenic threats) in relation to migration strategies and stopover choices between populations or individuals will be essential to help improve the current conservation and translocation efforts (see www.houbarafund.org).

